# Sample preparation protocol enabling nano-to-mesoscopic mapping of cellular connectomes and their habitats in human tissues and organs

**DOI:** 10.1101/533448

**Authors:** Lucy Ngo, Anton D. Nathanson, Tomasz Garbowski, Ulf Knothe, Dirk Zeidler, Melissa L. Knothe Tate

## Abstract

Multibeam scanning electron microscopy (multiSEM) provides a technical platform for seamless nano-to-mesoscale mapping of cells in human tissues and organs, which is a major new initiative of the U.S. National Institutes of Health. Developed for rapid throughput imaging of minute defects on semiconductor wafers, multiSEM has recently been adapted for imaging of human organs, their constituent tissues, and their respective cellular inhabitants. Through integration of geospatial approaches, statistical and network modelling, advances in computing and the management of immense datasets, as well as recent developments in machine learning that enable the automation of big data analyses, multiSEM and other cross-cutting imaging technologies have the potential to exert a profound impact on elucidation of disease mechanisms, translating to improvements in human health. Here we provide a protocol for acquisition and preparation of sample specimen sizes of diagnostic relevance for human anatomy and physiology. We discuss challenges and opportunities to integrate this approach with multibeam scanning electron microscopy work flows as well as multiple imaging modalities for mapping of organ and tissue structure and function.

## Introduction

*Connectomics* refers to the mapping of cellular connectivity in a physiological context, *e.g.* in organs and organ systems^1-3^. Seamless, multiscale and multimodal imaging provides a platform for interdisciplinary research and the trans-dimensional linking of biological structure and function ^1–11^. The U.S. National Institutes of Health’s (NIH) Connectome Project, launched in 2009, aimed to map “the human brain…to connect its structure to function and behavior”^10^. In 2018, NIH debuted a program to promote cross-cutting research mapping the cells within the human body^11^. Connectome maps of brain and other tissues of the body have typically been restricted to small organisms due to complications associated with cross-scale imaging from nano-to mesoscopic length scales. New imaging technology platforms enable such seamless scale-crossing image acquisition, and current barriers include biophysical hurdles intrinsic to specimen preparation as well as digital hurdles inherent to massive data management, analysis, and curation to maximize sharing across platforms and among geographically far flung users.

Electron microscopy (EM), a seminal technology in the advancement of cell biology and a foundational tool crossing disciplines, resolves subcellular detail and fine ultrastructure of cells. Currently, there is a great imperative not only to study the cells themselves but also to refocus at the intersection of and across multiple length scales encompassing the cellular networks that inhabit complex biosystems comprising tissues, organs, and organ systems. The status of these organ connectomes provides a powerful early indicator of degenerative changes and disease states over the lifespan of the organism. Three-dimensional multiscale imaging methods allow unprecedented study of structure and function across length scales; from organ to tissue, cells to molecules^7-9, 14^.

Traditional transmission electron microscopy (TEM), commonly used to visualise cellular ultrastructure, provides superior resolution and contrast. However, this is contingent on the often-laborious preparation of specimens into ultrathin sections (no more than 100 nm in thickness) for electron transparency. Standard EM specimen preparation techniques are optimised for small volumes, namely mouse-sized and smaller. Specimen sizes are constrained by diffusion path lengths of chemical fixation media as well as ultramicrotome chamber and diamond knife dimensions, not exceeding a few millimeters. Furthermore, image acquisition of tissue and organ-sized samples would take a matter of years with conventional EM^5^.

The emergence of multibeam scanning electron microscopy, with 61 or 91 parallel beams, is heralded by a new age of organ connectomics and presents a suitable platform for imaging large areas at nanometer resolution^1, 5-9^. Despite such pivotal advances in imaging technologies, biological tissue specimen preparation has remained unchanged and presents a barrier to translation of cutting-edge imaging modalities as well as to unleashing the potential of multimodal imaging. **This protocol addresses the difficulties associated with specimen preparation workflows enabling multibeam scanning electron microscopy (multiSEM) imaging as well as multimodal and correlative imaging methods.** While having been implemented successfully in peer-reviewed publications (Fig. 1), details of the protocol have been requested by the medical and scientific community alike, providing impetus for its publication.^5-7, 9, 15^

**Fig. 1:**
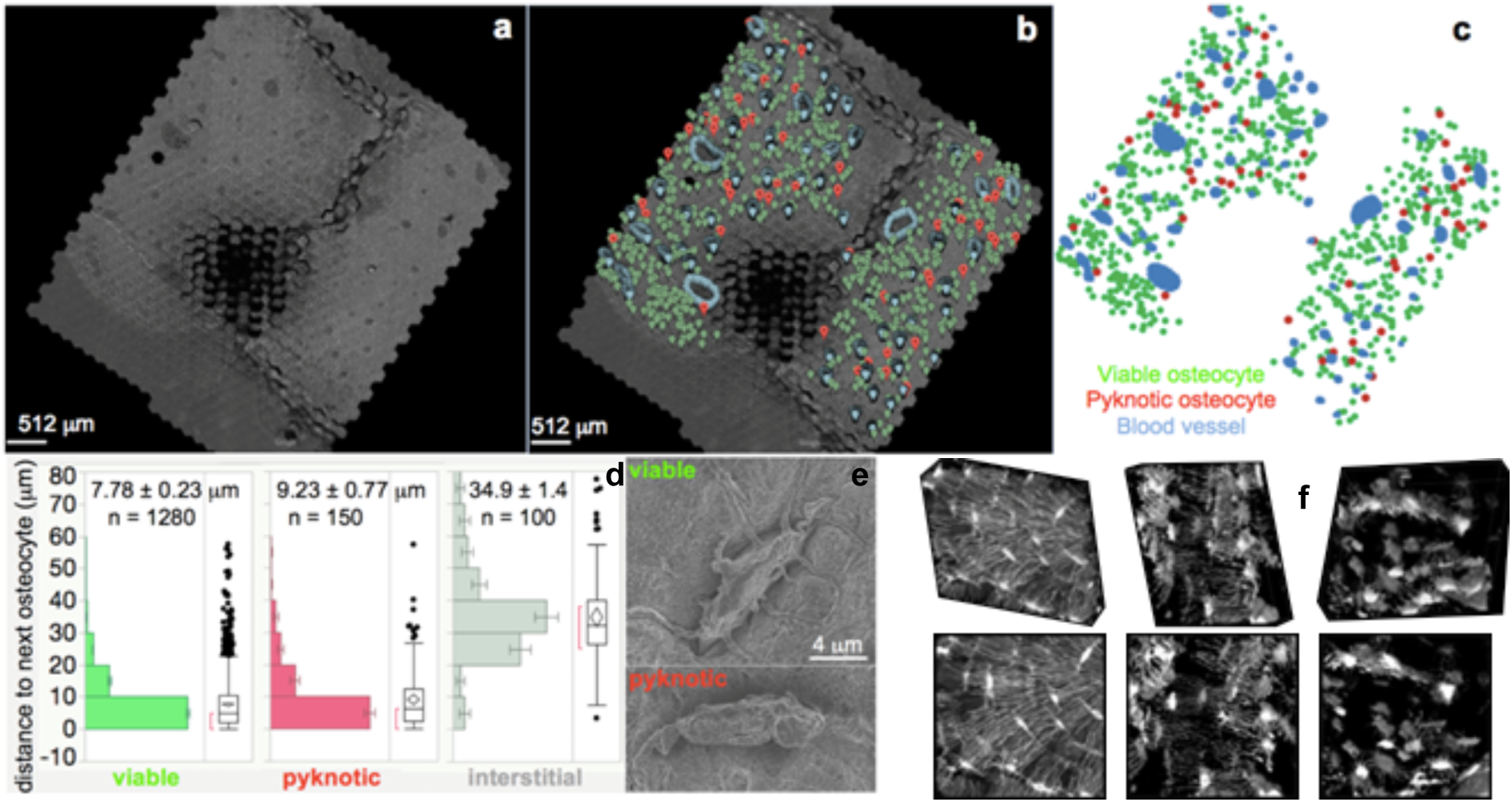
Cellular epidemiology studies in large scale maps comprising stitched, scanning electron microscopy images. These high resolution images are obtained using multibeam SEM and demonstrate the feasibility of the approach that greatly expands beyond fields of view previously acquired using confocal scanning microscopy. **a** Geonavigational methods and the Google Maps Javascript API were utilized to create navigable maps of osteocytes within the human hip imaged using multiSEM and spanning the nano-to mesoscale. The world’s first map of the human hip was created by imaging a sample approximately 2.4 × 2.4 mm. Imaging using a 61-parallel-beam Zeiss MultiSEM 505 prototype yielded over 55 000 SEM images, which were stitched into a single image, and rendered in a pyramidal structure for seamless zooming into and out of the map using Google Maps API. The dark area in the middle of the sample is an artifact of automated polishing, a sample preparation step that was later changed to precision CNC milling and hand polishing (see **detailed protocol**). **b, c** Google Maps Javascript API enabled the geographic coordinates of physiologically relevant landmarks to be recorded. Blood vessel edges, viable osteocytes and pyknotic (sick) osteocytes were marked with blue, green and red pins respectively. **d** The position of each pin could be exported for *post hoc* analyses, *e.g.* to test the hypothesis that cell viability correlates to blood vessel proximity, path lengths of viable, pyknotic (sick) and interstitial cells to blood vessels were calculated and compared. **e** MultiSEM images of a viable cell, with projections emanating outwards into the matrix and pyknotic cell, no projections observed. **f** The role of cell health and connectivity on tissue health was first observed in 3D images of osteocyte networks from (left) healthy, (middle) osteoporotic and (right) severely osteoporotic cortical bone from the femoral head of human patients, using laser scanning confocal microscopy. A shortcoming of this approach is that volumes of interest are limited to *e.g.* 274 × 274 × 50 microns. *Figures adapted and used with permission*^5-7, 9, 15^.

Presented in this protocol is a detailed workflow for sample preparation of sizeable tissue specimens (up to 10 cm in diameter, Fig. 2) of human and large mammalian origin. The method has been tested in human samples from the femoral neck and head of subjects undergoing hip replacement (note: all human samples collected and used in development of this protocol complied with all relevant ethical regulations of the Institutional Review Board of the Cleveland Clinic, IRB #12-335) as well from sheep (*Ovis aries*) femur and guinea pig (*Cavia porcellus*) knee to verify multimodal imaging (note: all animal tissue samples used in development of this protocol derived from studies that adhered to protocols approved through the respective Institutional Animal Care and Use Committees [IACUC] of the Canton of Grisons, Switzerland and Case Western Reserve University, in adherence with the Animal Research: Reporting of In Vivo Experiments [ARRIVE] guidelines). Here we adapted methods typically used for TEM and SEM tissue sample preparation to enable across length scales, from nano-to mesoscopic. The methods integrate techniques including precision CNC-milling and chemical etching methods developed for atomic force microscopy to enable three dimensional and multimodal imaging. This method can be applied to macroscopic human tissues and tissues of large mammals and can be integrated into a multimodal imaging workflow, in particular correlative light electron microscopy (CLEM).

**Fig. 2:**
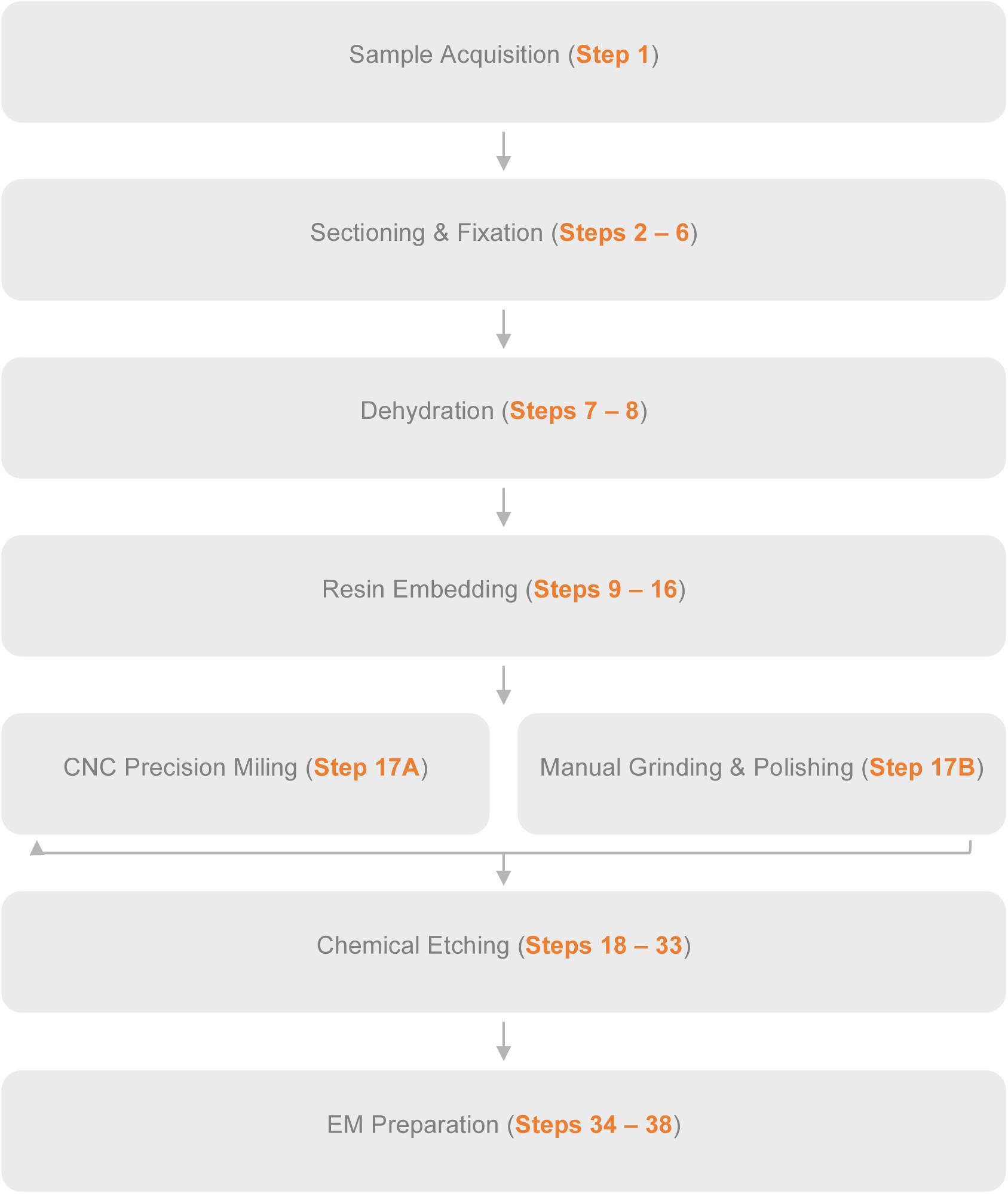
Workflow for acquisition of multiSEM data using mesoscopic samples.

**Fig. 3:**
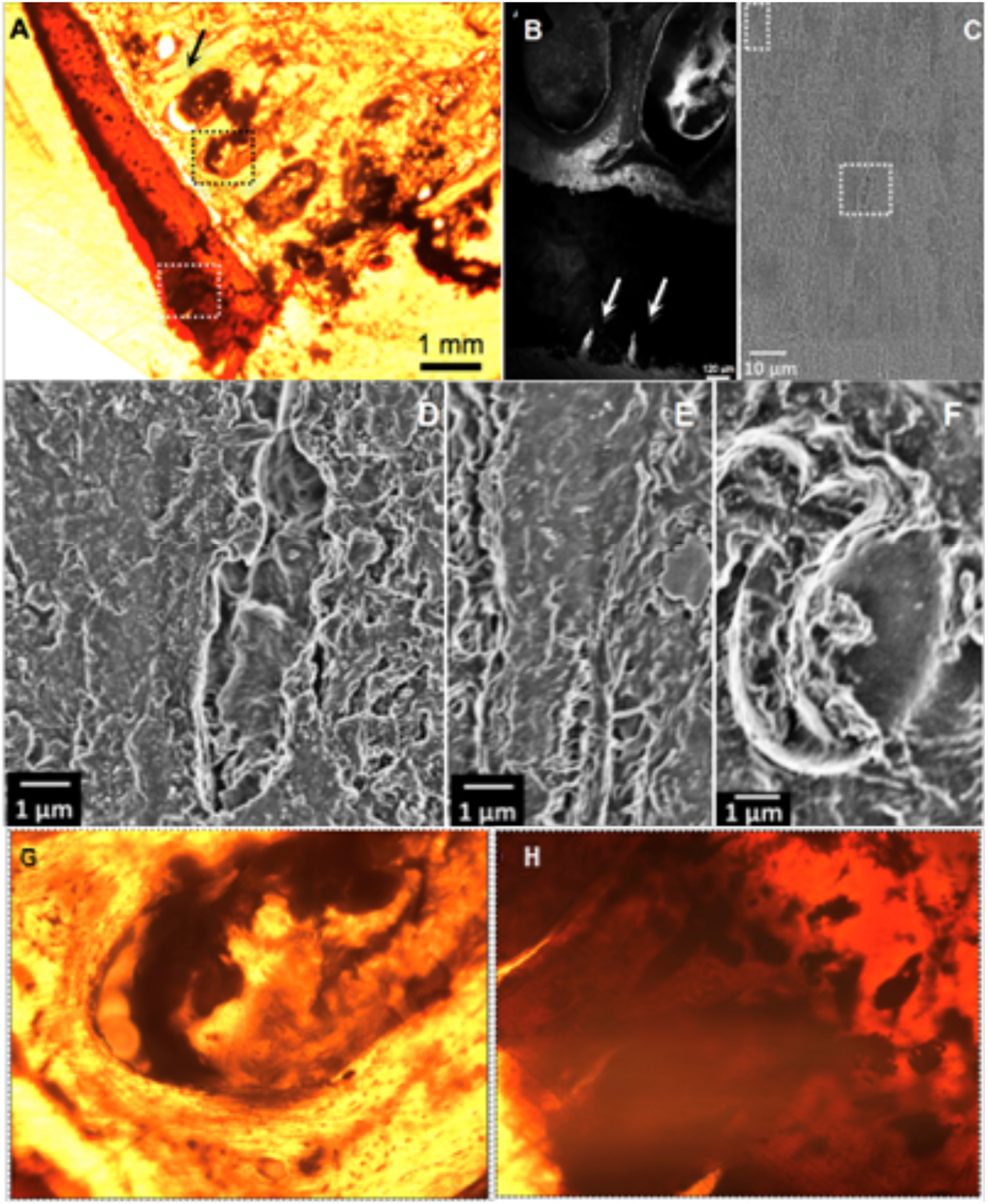
Seamless multi-scale imaging of hip joint tissue, prior to etching step. **A**, Transmitted light image of OsO_4_ stained femoral head section, with greatest penetration of contrast agent (darkest) in cartilage (white dotted square) and marrow spaces of the subchondral bone (black dotted square). **B**, Laser scanning confocal image of specimen shows autofluorescence (light areas) of subchondral bone and marrow spaces (ellipsoids) as well as cartilage (dark), with two visible cartilage defects (bright cracks, arrows). Scale bar 120 µm. **C**, Large scale, tiled multibeam SEM image of same area with single field of view showing (sub-)cellular resolution of bone (**D**, **E**) and blood (**F**) cells without additional etching. **G**, Subchondral bone exhibits less staining than cartilage and is highly cellular. Area corresponds approximately to black dotted square (in **A**). **H**, The tidemark, delineating the interface of unmineralized (rust-brown) and mineralized (orange) cartilage is visible in the upper right corner. Defects in the uncalcified cartilage are visible as light “cracks” running orthogonal to the tidemark line. Area corresponds approximately to the white dotted square (in A). Note regarding suboptimal quality of images: these images are taken of a 300 µm thick, EPON^®^ embedded, undecalcified section without prior mounting and cover slipping, using an inverted epifluorescent microscope with 1.6x (A) and 10x (**G**, **H**) dry objectives. Specimens were not mounted, or cover slipped due to plans to image subsequently using SEM. After completion of SEM studies, microradiography and renewed optical imaging can be carried out.

**Fig. 4:**
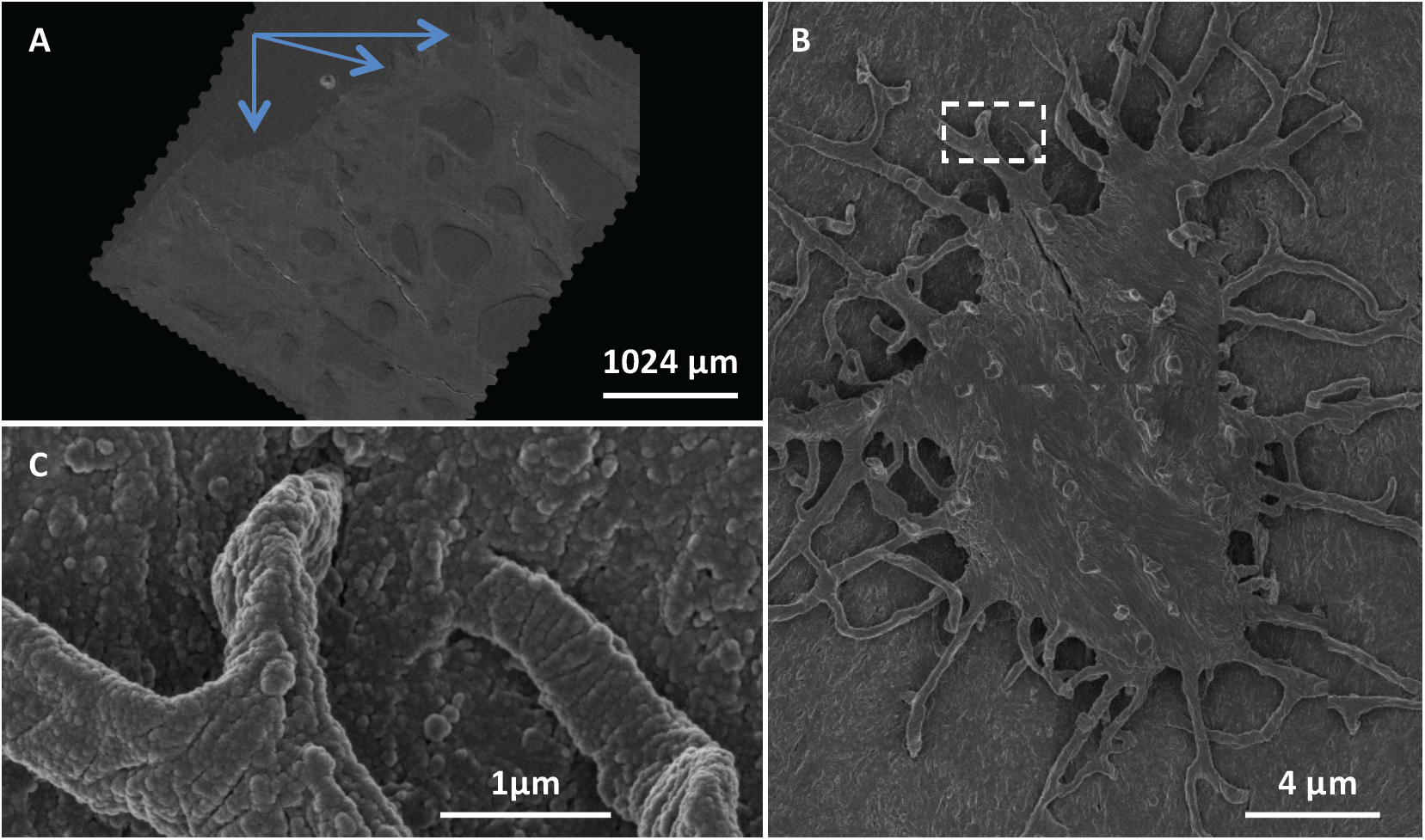
Seamless nano-to mesoscale imaging of the human hip, visualized using the Google Maps navigational platform. **A** The Google Maps JavaScript API was used to visualise and create navigable maps of an imaged region. The blue arrows point to the eroded surface of the hip joint, toward areas internal to the femoral head. **B** A single osteocyte on the sample surface with **C** projections (osteocyte processes) into the underlying matrix. This sample was imaged using a 61-parallel-beam Zeiss MultiSEM. The complete map can be accessed via http://www.mechbio.org/sites/mechbio/files/maps7/index.html [prior to publication, input of username and password are required to access the dataset]. username: mechbio password: #google-maps

This protocol is ideal for studies involving tissue samples from human or large mammals (up to 10 cm in diameter) where the organisation of cells and tissues are of primary interest. While the resultant data is of great detail, it is also of considerable size (>10TB) and requires a robust data management system or pipeline for efficient analytics and interpretation. The advent of deep learning and machine learning approaches is a keystone in the analysis of data sets of this size as manual methods would prove too costly and time consuming^16^.

## Comparison with other methods

The key advantages of this method for biological specimen preparation are that it is streamlined and flexible enough to be applied to any tissue specimen using readily available materials typically used in atomic force, EM and laser confocal microscopy specimen preparation. However, this flexibility and ease of implementation has its drawbacks as the method is not entirely self-contained and the sample must be moved from platform to platform if light microscopy is used in addition to EM. Advantageously, the use of heavy metal stains, uranyl acetate (UA) and lead citrate is eliminated as carbon sputtering provides sufficient contrast enhancement in the workflow. This further streamlines the process, avoiding the use of hazardous chemicals and their associated risks. Similarly, osmium tetroxide (OsO_4_) is not used in the fixation contrast as carbon sputtering alone provides sufficient contrast. This provides further advantages in context of multimodal, correlative microscopy, as OsO_4_ imparts a yellow hue throughout the specimen, impairing any form of light microscopy (Figs. 2-4).

The choice of polymethyl methacrylate (PMMA), over epoxy, results in samples robust and with little damage observed after electron beam exposure. MultiSEM requires sample flatness and parallelism to the mount surface to prevent charging artifacts, which ideally requires precision CNC-milling, an unconventional technique for EM sample preparation. Similar results have been achieved with conventional polishing methods.

The protocol does not involve ultra-thin sectioning characteristic of TEM, which is often difficult and time consuming. As a result use of the method may not perfectly preserve or enable visualization of internal cellular ultrastructure. For ultrastructure studies, TEM resolutions remain superior to EM. Neither the protocol method nor EM in general enable live-cell imaging, and images represent single points in time. Hence, the method does not capture the nuance and temporal dynamic of complex biological processes.

Novel techniques such as field emission scanning electron microscopy (FESEM)^17^ and serial block-face scanning electron microscopy (SBF-SEM)^18^ allow the three-dimensional visualisation of and reconstruction of cellular ultrastructure, however samples are limited to a few millimeters on edge in size. Although the images acquired from this protocol are two-dimensional, they provide a birds-eye-view, at nanometer resolution, of cellular networks and populations over large areas. The method thus enables visualization of tissue architecture not readily achievable using conventional two-dimensional EM.

### Experimental Design

For each experiment, we defined the relevant research question and a testable hypothesis, relating independent variables (also referred to as outcome measures) as a function of dependent variables. Exclusion criteria were typically defined at this stage to reduce effects of unrelated disease states that could confound study outcomes and interpretation, *e.g.* metabolic bone disease is excluded in a study of loss of bone density due to disuse.

Power calculations were made to determine sample size for each experimental group; although approximate, these accounted for variance in data based on pilot studies. Some particularly novel studies do not have a representative dataset upon which variance measures can be made and initial sample sizes have to be based on previous experience. Power analyses conducted for previous studies determined the need for a sample size of 5 patients *per* cohort. *Post hoc* analyses on these data indicated the robustness of study outcomes; from a translational perspective, even if more than 700 patients were tested to obtain statistical significance, differences (in periosteum derived stem cell regenerative capacity attributable to age and/or disease state) would remain small, due to the small to medium effect size (of age and disease), even if significantly different^19^.

### Specimen Acquisition

The critical phase of any research requiring the use of human tissue samples is the acquisition of specimens (Fig. 5). It is crucial to find a surgeon willing and able to collaborate in the study, acting as not only clinical investigator on the ethics protocol but ensuring the identification and screening of possible participants, alongside ensuring informed consent is obtained for each individual involved. It is crucial all specimens are de-identified, by removing all identifying information such as name, medical patient numbers, *etc.* by the tissue procurement and/or pathology department of the clinic prior to release of the tissues for scientific research. Age and sex of the patient are retained in sample documentation. Together with date of acquisition, specimens are separately identifiable without access to medical documentation.

**Fig. 5:**
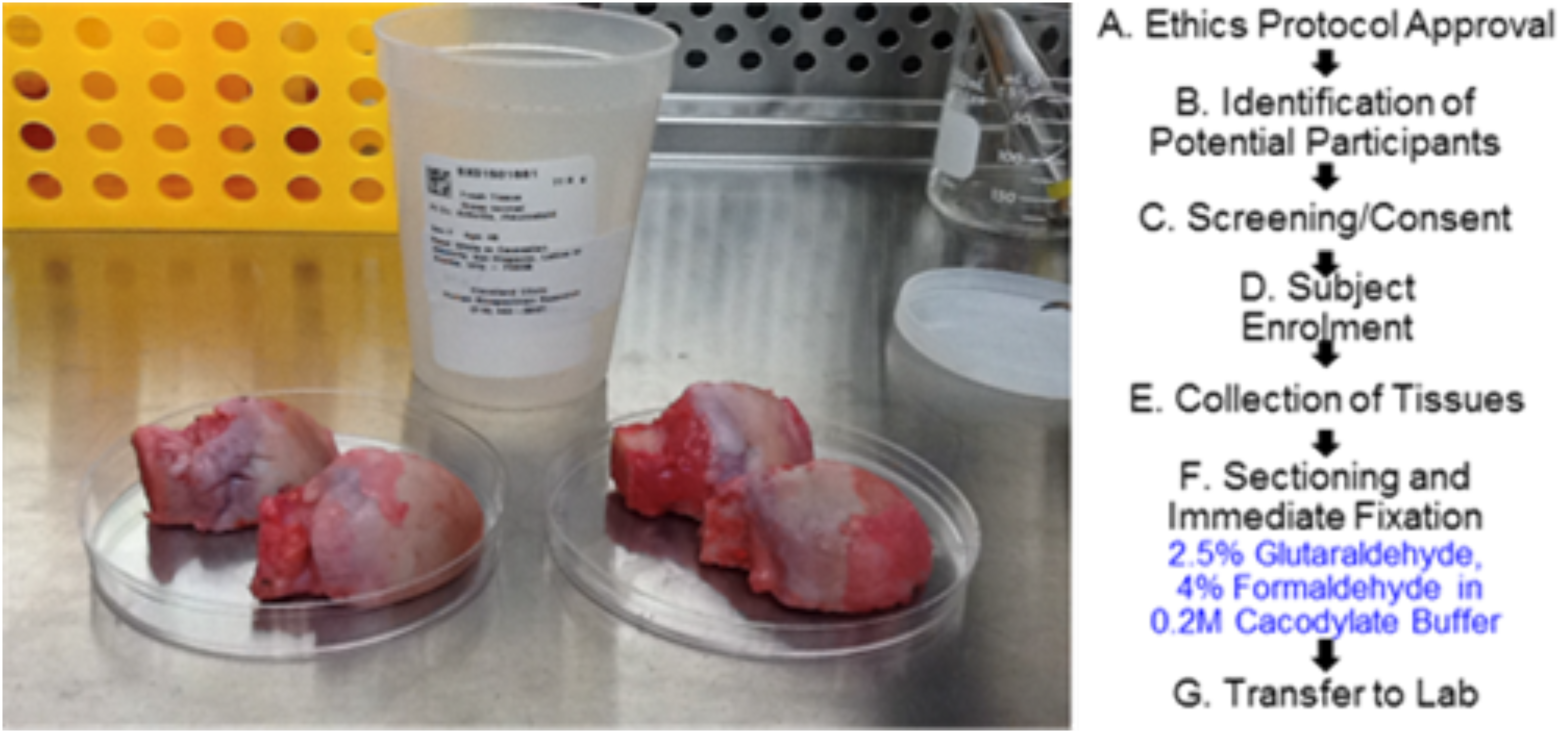
Procedures for acquiring and handling human tissues. Two samples from different patients, acquired on the same afternoon. Femoral heads (see photo, rounded structures divided into to halves) were sectioned along the plane of interest, *e.g.* coronal or sagittal plane, to allow for infiltration of fixation and embedding chemicals. **A-G** Work flow chart for sample acquisition, with chemicals indicated in blue.

In some experimental study designs, samples are given non-obvious identifiers and the key to decoding the identifiers is retained in a secure position by the clinical partner to enable later information from the patient’s medical record, *e.g.* specific health history information, if needed. In obtaining patient consent while also striving to include diverse populations in cohorts, consideration should be made to patient groups whose beliefs may preclude sharing of tissues. It may be possible to encourage their inclusion through awareness of cultural sensitivities and inclusion of language that addresses culture-specific considerations in outreach brochures as well as informed consent documentation.

### Sectioning and Fixation

With samples of a size larger than a millimeter on edge, it is important to take extra measures to ensure samples undergo adequate fixation before proceeding. In this protocol, we are limited to diffusion fixation; perfusion of fixatives is not an option in human tissues. The femoral head and neck are sectioned from the proximal and distal end before further sectioning of the medial and lateral portions, transversely across the bone. Following this, 1 mm slices in the plane of interest are taken, slices at this thickness allow the complete infiltration of glutaraldehyde and paraformaldehyde fixatives in cacodylate buffer. The specimens are then washed in cacodylate buffer before sequential ethanol dehydration (30%, 50%, 75%, 95%).

### Contrast Enhancement

Biological specimens for EM typically require post-fixation contrast enhancement to resolve subcellular detail. Contrast enhancing agents such as osmium tetroxide (OsO_4_), are commonly used but impart exogenous colour throughout the entire sample, making it non-conducive to multimodal imaging, namely light microscopy (Fig. 2). Heavy metal staining is another mode of contrast enhancement, where uranyl acetate (UA) and lead citrate are routinely employed. The use of heavy metals creates a more laborious and hazardous process, which was deemed neither desirable nor necessary. Additionally, UA introduces instability into the sample, making it light and UV sensitive and prone to forming precipitates over time. Instead we opted to highlight subcellular detail *post hoc*, with the use of both chemical etching (detailed below and adapted from protocols used for AFM), and carbon sputtering.

### Resin Embedding

Conventionally, biological specimens are embedded in epoxy resins for EM. Initial attempts at using epoxy, EPON^®^, proved unsuccessful as it is typically used for samples that are 1-2mm on edge. Under inspection with a light microscope, the epoxy embedded specimens exhibited variations in surface topography of up to 100µm making it unsuitable for multiSEM. The specimens exhibited incomplete and non-uniform resin polymerization, most likely caused by insufficient resin penetration in such large specimens. Additional contributing factors to poor polymerization, include insufficient time for immersion and perfusion, inadequate dilution steps and excessive resin viscosity, all resulting in poor resin infiltration into tissue pores, leaving residual ethanol and propylene oxide within the sample.

Due to the difficulties encountered with EPON^®^ and prior experience with PMMA, we used lower viscosity polymethylmethacrylate (PMMA) resin as the embedding medium. To ensure complete infiltration of the resin into tissue pores and uniformity of the sample surface, dehydration time was increased, and the tissues were bulk embedded under vacuum to enable gradual polymerization.

### Planarity and Sample Size

The development of multiSEM enables the rapid throughput of specimens that far exceed traditional dimensions, up to 100 × 100 × 30 mm. However, multiSEM requires specimens to be flat and parallel to the stage, with a surface flatness less than or equal to 500 nm *per* 100 µm (peak-to-peak). This minimizes edge effects and specimen damage resultant of increased electron emissions at high points in the surface topography.

Initially, specimens were manually and automatically mechanically ground and polished, with both methods exhibiting various degrees of success. Automated polishing proved too aggressive, rapidly removing the sample surface. Conversely, manual polishing was not time effective and resulted in the emergence of microscopic cracks in the sample surface, most likely due to the release of residual stresses. These cracks could later one cause edge effects along the edges of the crack, interfering with EM imaging quality. We progressed to micro-CNC milling and manual buffing with acrylic polish, the sample surface was milled with a great level of precision without the introduction of surface cracks. Specimens were precision milled in a CNC-lathe using a carbide insert with the unhoned edge against the sample surface. Following this, the surface was lightly buffed with a soft microfiber cloth, acrylic polish and ultra-pure water.

### Chemical Etching and EM Preparation

As noted above, this protocol does not use post-fixation contrast enhancement with heavy metals. Instead, chemical etching methods adapted from techniques developed for AFM were used to remove uncover underlying molecular detail and cellular networks^20, 21^. Dilute hydrochloric acid (HCl) and sodium hypochlorite (NaClO) enable controlled etching of the organic, collagen, and inorganic, apatite composite matrices, revealing cells and projections embedded in the extracellular matrix. This method also allows for selective etching of regions of interest by masking around areas to be etched with electrical tape (geometric shapes) or nail polish (free form regions). Following this step, the sample can be imaged with light microscopy. To increase sample conductance and contrast, the specimen requires a thin carbon coat (20 nm) before it can be EM imaged.

## Materials

### Reagents

▪ Milli-Q ultra-purified water (18.2 MΩ cm, Millipore Milli-Q Reference Water Purification System, cat no. Z00QSV001)
▪ Tissue Samples for fixation ! CAUTION Tissues should be harvested according to relevant ethics protocols.

### Immersion fixation

▪ Glutaraldehyde, 10% aqueous solution (Electron Microscopy Sciences, cat no. 16100) ! CAUTION It is hazardous upon inhalation and skin contact. Wear gloves, protective clothing and eyewear, and use under a fume hood.
▪ Paraformaldehyde Powder (Sigma-Alderich, CAS. 30525-89-4) ! CAUTION It is hazardous upon inhalation and, skin and eye contact. Wear gloves, protective clothing and eyewear, and use under a fume hood.
▪ Sodium cacodylate buffer (0.2 M, pH 7.4; Electron Microscopy Sciences, cat. no. 11650) ! CAUTION It is toxic upon ingestion and eye contact, wear gloves and protective eyewear.
▪ Aluminium foil (Sigma-Aldrich, cat. no. Z691569-1EA)

### Dehydration and polymer infiltration

▪ Ethyl alcohol series (ethanol, Rossville Gold Shield, cat. no. 412804, series 50, 70, 95, 100% (vol/vol)) ! CAUTION It is hazardous upon skin and eye contact, and flammable. Wear gloves, protective clothing and eyewear. ! CAUTION It is hazardous upon ingestion, skin and eye contact, and is flammable. Wear gloves, protective clothing and eyewear.
▪ Sodium Hydroxide (Sigma-Alderich, CAS. 1310-73-2) ! CAUTION It is toxic upon inhalation and, skin and eye contact. Wear gloves, protective clothing and eyewear, and use under a fume hood. Methyl methacrylate contains hydroquinone as stabilizer (Sigma-Alderich, CAS. 1310-73-2)
▪ Methyl methacrylate monomer, stabilised (CAS Number 55935-46-1) ! CAUTION Cleaned monomer is highly flammable and explosive and must be stored in an explosion proof refrigerator at 4°C prior to use.
▪ PMMA pellets (Sigma-Alderich, CAS. 55935-46-1)
▪ Calcium chloride anhydrous (pellets, 4-20 mesh) (Fisher Scientific, CAS. 10043-52-4) ! CAUTION It is hazardous upon eye contact, and flammable. Wear gloves and protective eyewear.
▪ Benzoyl peroxide powder ! CAUTION It is hazardous upon eye contact and is highly flammable and explosive. Wear gloves and protective eyewear

### Chemical Etching

▪ Hydrochloric Acid 5M (HCl, Lowy Solutions, cat. no. UCS-BIO-0035) ! CAUTION It is hazardous upon skin and eye contact, inhalation and ingestion. Wear gloves, protective clothing and eyewear.
▪ Sodium Hypochlorite 12.5% (NaClO, Chem-Supply Pty Ltd Australia, CAS. 7681-52-9) ! CAUTION It is hazardous upon inhalation and, skin and eye contact. Wear gloves, protective clothing and eyewear, and use under a fume hood.
▪ Methanol (Electron Microscopy Sciences, cat. no. 18510) ! CAUTION It is hazardous upon skin and eye contact, inflammation and is highly flammable. Wear gloves, protective clothing and eyewear.
▪ Ethanol, Absolute (Sigma-Alderich, CAS. 64-17-5) ! CAUTION It is an irritant upon skin and eye contact, and flammable. Wear gloves, protective clothing and eyewear

### Equipment

▪ Nitrile gloves (Livingston item. no. GLVNTRLC100S)
▪ Dust-free wipes (Professional Kimtech Science Kimwipes Delicate Task Wipers, Kimberly-Clark Professional, no. 34155)

### Sample Sectioning

▪ Bandsaw (Craftsman)
▪ Precision Cutter (Isomet 4000 Linear Precision Saw, Buehler)
▪ Saw Microtome (Leica SP1600 Saw Microtome, Leica Biosystems)

### Immersion fixation

▪ Plastic Petri dishes (100mm)
▪ Parafilm M (Fisher Scientific, cat. no. 13-374-10)
▪ Forceps

### Embedding and Polymerization

▪ Separatory Funnel
▪ Ring stand
▪ Buchner Funnel (lined with filter paper)
▪ 500ml Glass jars with lids
▪ Mallet

### Surface Preparation

▪ CNC Lathe with Carbide Insert without a honed edge
▪ Acrylic Polish (Novus Fine Scratch Remover 2)
▪ Soft Microfiber Cloth
▪ Semiautomatic Grinder and Polisher (Struers LaboPol-5)
▪ 1200-grit Grinding Disk (Struers MD-Piano 1200)
▪ 500-grit Grinding Disk (Struers MD-Piano 500)
▪ Polishing Disk for 3 µm suspension (Struers MD-Dac)
▪ Polishing Disk for 1 µm suspension (Struers MD-Nap)
▪ 3 µm Diamond Suspension Polishing Compound (Struers DiaPro Dac 3 µm)
▪ 1 µm Diamond Suspension Polishing Compound (Struers DiaPro Nap 1 µm)

### Chemical Etching and EM Preparation

▪ Transfer Pipettes (Thermofisher Scientific, Catalogue number 202-1S)
▪ Glass Petri Dishes (80mm diameter)
▪ Glass Crystallising dishes (125mm diameter)
▪ Lint free cotton swab with anti-static handle (Puritan Medical Products, SKU#: 876-PPC)
▪ Evaporative Carbon Coater (JEE-420 Evaporative Coater, JEOL)
▪ Silver Paste (Electron Microscopy Sciences, cat no. 12640)
▪ Ultrasonic Cleaner (Digitech)

## Reagent Setup

### 16% (wt/vol) paraformaldehyde solution

▪ To make the paraformaldehyde solution, add 8g of paraformaldehyde into 50ml of ultra-pure H_2_O. In a fume hood, stir the mixture at 60°C until fully dissolved. Cover with aluminium foil and allow to cool., add 2g paraformaldehyde to 40ml dH_2_O. Prepare fresh prior to use.

### 2.5% (vol/vol) glutaraldehyde and 4% (wt/vol) paraformaldehyde in 0.1 M cacodylate buffer

▪ To make 200ml, mix 50ml of 10% (vol/vol) glutaraldehyde with 50 ml of 16% (wt/vol) paraformaldehyde in 100 ml of 0.2 M cacodylate buffer. Prepare fresh prior to use.

### 0.02M Hydrochloric Acid Solution

▪ To make 200ml, add 800µl of 5M hydrochloric acid and make up the volume to 200ml using ultra-pure H_2_O

### Sodium Hypochlorite Solution

▪ To make 200ml, mix 160ml of 12.5% sodium hypochlorite with 40ml of ultra-pure H_2_O.

### Cleaned Monomer

▪ Mix 1000ml of 5% NaOH and 1000ml of methyl methacrylate (MMA). Allow the solutions to separate, then drain off only the lower solution for disposal, repeat twice. Then add 1000ml dH_2_O to the remaining solution, allow it to separate and drain off the lower solution for disposal, repeat twice. Drain the MMA into a flask through a Buchner funnel lined with filter paper and contains 100g of calcium chloride pellets. Repeat with new calcium chloride pellets. Allow monomer to reach room temperature before use (25°C). ▲ CRITICAL STEP The methyl methacrylate monomer contains inhibitor hydroquinone to prevent polymerization in transit. The hydroquinone must be removed prior to use. To do this it is ‘washed’ with 5% sodium hydroxide solution. ! CAUTION Cleaned monomer is highly flammable and explosive and must be stored in an explosion proof refrigerator at 4°C prior to use.
▪ PAUSE POINT Methyl methacrylate monomer can be stored for six months in an explosion proof refrigerator at 4°C.

### Polymer preparation

▪ Add 3g of dried benzoyl peroxide to 300ml of clean monomer in the glass screw-top jar and stir until dissolved. Add 120g methyl methacrylate beads to the jar gradually over a period of 2-3d, stirring constantly and allowing beads to dissolve completely between additions.

## Procedure

### Sample Sectioning and Fixation • TIMING 11d

! CAUTION To avoid exposure to hazardous substances, all steps should be performed in a chemical fume hood with appropriate protective gear (gloves, lab coat and eye protection).

1. Femoral necks and head samples are sourced from patients who have undergone hip replacement surgery. ! CAUTION Particular care needs to be taken in handling of human tissues, which carry a risk of disease transmission via blood borne pathogens if inadequate safety procedures (prevention of contact through use of gloves, safety glasses, laboratory coats, and biohazard safety cabinets) are taken. Also, during transport from the clinic to the lab, extra care must be taken to ensure that the sample remains completely contained, e.g. through double bagging of the sample in containers contained in zip-locked specimen bags. ▲ CRITICAL STEP Use of human samples requires Institutional Review Board (IRB) approval, also referred to in some countries as Human Ethics Protocols. Use of tissues and organ samples from cadaveric donors also requires ethics review board approval.
2. Cut the femoral head into three sections from proximal end to distal end.
3. Cut the medial and lateral portions into three pieces transversely across the bone.
4. Section tissue into 1mm thick section along the plane of interest e.g. coronal or sagittal plane, to allow for infiltration of fixation and embedding chemicals
5. Fix the sectioned tissue immediately after with 2.5% glutaraldehyde, 4% paraformaldehyde in 0.2M cacodylate buffer at room temperature for 5 d ! CAUTION Fixative is hazardous. ! CAUTION Samples should be collected, sectioned, and fixed as soon after resection as possible to preserve sample properties.
6. Wash sample with cacodylate buffer (0.2M, pH7.3) daily for 5 d
  ▪ PAUSE POINT Samples can be stored in cacodylate buffer for up to 2 w.

### Dehydration • TIMING 9 d

7. Dehydrate the sample in an ethanol series (30%, 50%, 75%, 95%) for 3 d each, changing daily
8. Place the sample in absolute ethanol for 3 d, changing daily

### Embedding and Polymerization • TIMING 2 – 10d

9. Embed sample in polymer by filling glass container with polymer.
10. Insert specimen with section side facing down.
11. Cover with foil and place in explosion-proof refrigerator for 48 h
12. Periodically remove from refrigerator, checking orientation.
13. Allow container to remain at room temperature (25°) until polymerization occurs (3-7 d). Once polymerization extends slightly above the specimen, checking hardness with a teasing needle, place into a 60°C incubator overnight to complete polymerization.
14. Place specimen in plastic bag and strike the glass with a mallet to shatter it.
15. Carefully remove block and rinse under running H_2_O.
16. Trim excess plastic from block with bandsaw
  ▪ PAUSE POINT Samples can remain in storage indefinitely before proceeding to the following methods.

### Surface Preparation • TIMING 0.5 – 1h

1. Choose option A if precision-CNC milling is available. Choose option B if precision CNC milling is not possible but semi-automatic and hand polishing is an option.
  A. Precision CNC-milling of bulk embedded samples
    i. Fix the sample to a CNC-lathe
    ii. Mill to require depth using a carbide insert without a honed edged facing the sample surface
    iii. Gently polish the sample surface with a soft microfiber cloth, acrylic polish and ultra-pure H_2_O.
    iv. Flow ultra-pure H_2_O over the sample to clear of all polish and residue.
    v. Gently clean the sample with absolute ethanol and a lint free wipe
    vi. Ensure the sample is thoroughly dry and free of dust by placing it under a steady stream of inert gas (nitrogen).
  B. Manual Grinding and Polishing
    i. Mount the 500-grit polishing disc on the polishing wheel
    ii. Start the polishing wheel at 200rpm and place the surface face down with even pressure
    iii. Gradually resurface the sample using the 500-grit then lift from the wheel.
    iv. Place the sample in a beaker of H_2_O and into an ultrasonic cleaner.
    v. Stop the wheel and then mount the 1200-grit polishing disc
    vi. Start the polishing wheel at 200rpm and place the surface face down with even pressure
    vii. Gradually resurface the sample using the 1200-grit then lift from the wheel.
    viii. Place the sample in a beaker of H_2_O and into an ultrasonic cleaner.
    ix. Stop the wheel and mount the Dac polishing pad and apply a small amount of 3 µm diamond suspension.
    x. Turn the polishing wheel to 200rpm and spread the suspension across the polishing pad.
    xi. Press the sample face down with even pressure and gently polish then lift from the wheel.
    xii. Place the sample in a beaker of H_2_O and into an ultrasonic cleaner.
    xiii. Stop the wheel and mount the Nap polishing pad and apply a small amount of 1 µm diamond suspension.
    xiv. Turn the polishing wheel to 200rpm and spread the suspension across the polishing pad
    xv. Press the sample face down with even pressure and gently polish then lift from the wheel.
    xvi. Place the sample in a beaker of H_2_O and into an ultrasonic cleaner.
    xvii. Gently clean the sample with absolute ethanol and a lint free wipe
    xviii. Ensure the sample is thoroughly dry and free of dust by placing it under a steady stream of inert gas (nitrogen).

▪ PAUSE POINT Samples can remain in storage indefinitely before proceeding to the following methods.

### Chemical Etching • TIMING 10 – 30 m

! CAUTION To avoid exposure to hazardous substances, all steps should be performed in a chemical fume hood with appropriate protective gear (gloves, lab coat and eye protection).

! CAUTION Gloves must be worn at all times during sample preparation and transfer. Do not touch the samples, sample holders or any parts or tools coming in contact with the sample with bare hands. Oil and grease from hands introduce contaminants to the SEM system and may cause damage.

18. Mount the sample onto the specimen holder, fixing it in place using silver paint (alternately carbon or copper tape may be used).
19. Gently clean the sample with absolute ethanol and a lint free wipe
20. Ensure the sample is thoroughly dry and free of dust by placing it under a steady stream of inert gas (nitrogen).
21. Create edges of etch zone and protect areas of surface not to be etched by covering a large margin using a masking medium i.e. electrical tape and/or finger nail polish ▲ CRITICAL STEP The masking medium acts as a resist to protect areas from the etching reagents. Ensure that a strong seal or bond is created between the electrical tape or nail polish and the sample.
22. Pipette off 2 ml of HCl solution and place on the surface of the sample that is to be etched ▲ CRITICAL STEP HCl solution should be enough to cover the sample surface, increase volume as required.
23. Gently expose entire surface of etch area to solution using a cotton swab for 60 s.
24. Rinse with copious ultrapure H_2_O and blot dry with dust-free wipes.
25. Pipette off 2 ml of HCl solution and place on the surface of the sample that is to be etched ▲ CRITICAL STEP HCl solution should be enough to cover the sample surface, increase volume as required.
26. Gently expose entire surface of etch area to solution using a cotton swab for 25 s.
27. Rinse with copious ultrapure H_2_O and blot dry with dust-free wipes.
28. To remove flaked off organic material, pipette off 2 ml of NaClO and place on the surface of the sample that is to be etched. ▲ CRITICAL STEP NaClO solution should be enough to cover the sample surface, increase volume as required.
29. Gently expose entire surface of etch area to solution using a cotton swab for 60 s.
30. Rinse with copious ultrapure H_2_O and blot dry with absorbent, lint free wipes.
31. Gently clean the sample with absolute ethanol and a lint free wipe and if electrical tape has been removed, making sure to remove any adhesive residue.
32. Ensure the sample is thoroughly dry and free of dust by placing it under a steady stream of inert gas (nitrogen). ! CAUTION Residual oil and dust introduce contaminants to the SEM system and may cause damage. Dust on the sample may hinder imaging.
33. Place in a protective specimen storage box to prevent exposure to elements or dust prior to carbon sputtering and EM imaging. ! CAUTION Storing samples for prolonged periods of time may lead to dust and other contaminants settling on the sample surface.

▪ PAUSE POINT Samples can remain in storage indefinitely before proceeding to the following methods. At this point samples can also be taken to be imaged with light microscopy.

### Preparation of sample for imaging • TIMING 0.5 – 1h

34. Gently clean the sample with absolute ethanol and a dust-free wipe.
35. Ensure the sample is thoroughly dry and free of dust by placing it under a steady stream of inert gas (nitrogen).
36. Carbon-coat the sample with a 20nm thick layer of film
37. Mount the specimen to the stage or metal stub using silver paint (alternately, carbon or copper tape can be used), ensuring an unbroken and robust connection is made between the sample surface and sample holder. ▲ CRITICAL STEP If carbon or copper tape is used, ensure there are no air bubbles or gaps between the tape and the specimen by rubbing and burnishing the tape against the specimen.
38. Place the sample under vacuum in preparation for imaging in accordance to your electron microscopy system.

## Acknowledgements

This work has been supported in part by the Paul Trainor Foundation and the Alexander von Humboldt Foundation. We would like to acknowledge Gregor Dellemann of Zeiss for his support of the project since its inception in 2011, Professor Thomas Bauer (Orthopaedic Pathology, Cleveland Clinic) for his expert advice and assistance, as well as Dr. Judy Drazba and her team at the Core Research Services of the Lerner Research Institute (Cleveland Clinic Foundation), including Mei Yin and Diana Mahovic, for their assistance with embedding of the specimens in EPON^®^ and PMMA. We would also like to acknowledge Dr. Nicholas Ariotti and Natasha Kapoor-Kaushik for technical assistance and use of facilities supported by AMMRF at the Electron Microscope Unit at UNSW. The contributions of Paul Feng, a cooperative student from Institut Polytechnique de Grenoble (Grenoble INP) who assisted with map generation, are also acknowledged with appreciation.

